# Temporal dynamics of implicit memory underlying serial dependence

**DOI:** 10.1101/2020.11.17.386821

**Authors:** Cristiano Moraes Bilacchi, Esaú Ventura Pupo Sirius, André Mascioli Cravo, Raymundo Machado de Azevedo Neto

## Abstract

Serial dependence is the effect in which the immediately preceding trial influences participants' responses to the current stimulus. But for how long does this bias last in the absence of interference from other stimuli? Here, we had 20 healthy young adult participants (12 women) perform a coincident timing task using different inter-trial intervals to characterize the serial dependence effect as the time between trials increases. Our results show that serial dependence abruptly decreases from 0.1 s to 1 s inter-trial interval, but it remains pronounced after that for up to 8 s. In addition, participants' response variability slightly decreases over longer intervals. We discuss these results in light of recent models suggesting that serial dependence might rely on a short-term memory trace kept through changes in synaptic weights, which might explain its long duration and apparent stability over time.

---

Temporal dynamics of implicit memory underlying serial dependence Perception and action can be biased due to the characteristics of recently presented stimuli. This serial dependence effect has been shown in the visual domain for a wide range of features, from low-level orientation (e.g., Liberman et al. 2016, Fischer and Whitney 2014) and speed perception (Makin et al. 2008; Kwon and Knill, 2013), to high-level face perception (Liberman et al. 2014). To exert its effect on the current trial, information from the previous trial's stimulus should be stored in memory. However, it is unclear how long this memory trace lasts.

Previous investigations on the temporal dynamics of stimulus information kept in short-term memory have shown that it can be stored with good levels of detail for up to 30 s (Magnussen & Greenlee 1992; Blake 1997), but that its precision decreases over time (Rademaker et al. 2018; Shin et al. 2017) or its representation drifts (Schneegans and Bays, 2018; Wolff et al. 2020). However, these studies have investigated short-term memory effects when keeping stimulus information in memory was both task-relevant and explicit. On the other hand, serial dependence emerges even when information from previous trials is task-irrelevant and does not need to be explicitly encoded in short-term memory (Collins 2020; Lieberman et al. 2016), suggesting that this bias does not depend on the active maintenance of information in memory on the previous trial. Critically, the temporal dynamics of such implicit memory traces are still mostly unknown.

Recent studies have shown that events from up to 4 trials into the past can modulate performance in the current trial (Kalm and Norris, 2018; Fischer and Whitney, 2014). Nevertheless, these experiments could not evaluate the pure progress of memory, given that the serial dependence effect was evaluated over intervening trials, which might have overwritten memory traces (Kalm and Norris 2018; Lewandowsky, Oberauer, Brown, 2009). When investigating the sole effect of time on serial dependence, Bliss et al. (2017) verified that after 10 s the previous stimulus would no longer bias current responses. In contrast, Papadimitrou et al. (2015) have shown that serial dependence persists for up to 6 s intervals. In both these studies, participants had to explicitly encode the current trial stimuli in short-term memory to perform the task. Therefore, it remains to be shown how serial dependence changes for varying inter-trial intervals without explicitly encoding stimulus information in short-term memory and without intervening trials. To characterize the effect and temporal dynamics of implicitly encoded information on serial dependence, we had participants perform a coincident timing task, in which they only had to act upon the current stimulus, without requiring them to encode items in short-term memory explicitly on the previous trial.

## Methods

### Participants

Twenty participants (12 women, 21±4 years old, mean ± standard deviation) were informed about the experimental procedures and signed a Consent Form. We recruited a convenience sample of college students. The experimental protocol was approved by The Research Ethics Committee of the Federal University of ABC. All experiments were performed following the approved guidelines and regulations. All participants had normal or corrected to normal vision.

### Stimulus and task

Participants performed a coincident timing task in which they pressed a button when a rightward moving target hit an interception zone (Figure 1A). The beginning of each trial was cued by presenting the start zone (squared black letter “C”) on the left side of the screen and the interception zone (vertical black line fitting 30% of the screen vertically), separated by 10 dva. Simultaneously, the target (red square, 0.5 degrees of visual angle [dva]) was flashed for 200 ms to warn participants the trial was about to begin. After 400 ms, the target was presented and started moving. Participants were instructed to press a button at the same time the target hit the interception zone (button box sampling frequency 1000 Hz). After 500 ms of target arrival, the target, start and interception zones were removed from the screen. Targets moved at a constant speed of 20, 22, 24, 26, or 28 dva/s on a gray background and took approximately 500, 454, 416, 384, and 357 ms to travel across the screen, respectively. To perform the task, participants sat in a quiet, dark room, at a distance of approximately 65 cm from a ViewPixx display monitor (120 Hz refresh rate, resolution of 1920 × 1080 pixels), resting their heads on a chin rest. Stimuli and task were generated and controlled using GNU Octave (https://www.gnu.org/software/octave/) and Psychtoolbox (Brainard, 1997; Pelli, 1997; Kleiner et al., 2007), running on a Linux operating system. Throughout the experiment, participants were instructed to fixate their gaze at a cross (0.5 dva) on the center of the screen, 0.35 dva below the target’s path. The fixation cross was present on the screen for the entire experiment.

**Figure 1.**
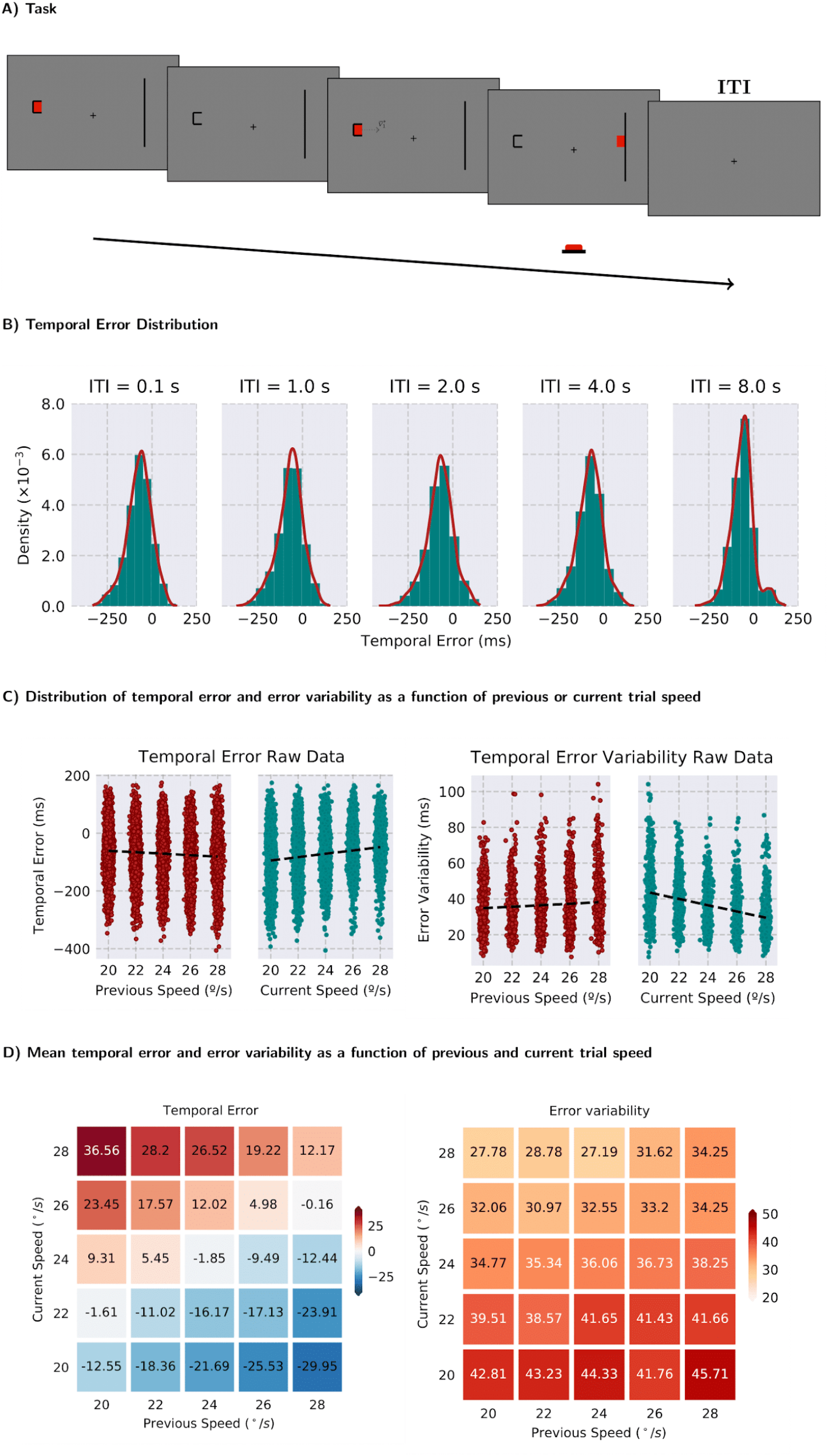
(A) Phases of a trial in the experiment. Participants were instructed to press the response button at the same time the target reaches the vertical bar. (B) Temporal error density for all participants collapsed by ITI. Values centered at zero and ranging between −200 and 200 milliseconds indicate that participants were performing a coincident timing task accordingly and not waiting for the target to reach the vertical bar to press the button. (C) Distribution of temporal error and error variability across participants as a function of previous (red) and current (blue) target speeds. Despite variability across participants, temporal errors tend to be more negative when the previous trial is faster and more positive when the current trial is faster. The error variability follows an opposite pattern: the greater the current trial speed, the lower the error variability across trials, and the lower the previous trial speed, the lower the error variability across trials. (D) Mean temporal error (left) and error variability (right) for all volunteers as a function of previous (horizontal axis) and current (vertical axis) target speed. The temporal error becomes more positive as the current target is faster (cells get redder from bottom to top), but also becomes more negative as the previous speed is faster (cells get bluer from left to right). The error variability shows the opposite pattern from temporal error.

### Experimental design and procedures

Before the main experiment, participants were instructed and familiarized with the task by performing 20 trials. Each target speed was randomly presented four times during practice. To evaluate the dynamics of the serial dependence effect, we had participants perform the coincident timing task in five separate blocks with different inter-trial intervals (ITI): 0.1, 1, 2, 4, and 8 s. The ITI was defined as the interval between the disappearance of the start and interception zones and its reappearance on the next trial.

Each block consisted of 5 mini-blocks containing 51 trials each. Within each mini-block, the target speeds were counterbalanced, such that each target speed was preceded by every other speed the same number of trials (Brooks, J.L., 2012). This sequence of trials assured that the current and previous target speed were independent of each other. Participants had 30 s rest periods between mini-blocks. Participants were allowed to take longer breaks between blocks. The experimental session took approximately 2 hours and comprised a total of 1275 trials.

### Analysis

We calculated participants’ temporal error (in ms) on each trial, i.e., the temporal difference between the time participants pressed the button and the target arrival at the interception zone. Negative values indicate early responses, whereas positive values indicate delayed responses. In addition to the temporal error, we also estimated participants’ error variability by calculating the standard deviation of participants’ temporal error on every combination of previous and current trial speeds for each ITI.

We first removed outlier trials for each ITI and participant by calculating the median absolute deviation (MAD) and excluding all trials where the deviation was greater than 3 (Leys et al., 2013; Rousseeuw et al., 1993). We removed one participant from the 8 s ITI condition because more than 15% of trials were removed. Overall, 2.81% of trials were removed with this procedure.

We evaluated serial dependence by separately modeling participants' temporal error for each ITI using multiple linear regression,

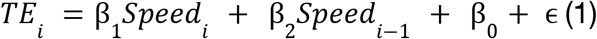

where *TE*_*i*_ is the temporal error (or the variability of the error) on trial *i*, β_1_ is the slope of the speed on the current trial and β_2_ is the slope of the speed on the previous trial, β_0_ is the intercept and ϵis a Gaussian error centered at zero. The main measure of serial dependence was β_2_, the slope of the previous trial speed on the multiple linear regression. It is important to note that, like in other studies evaluating serial dependence using different tasks (e.g., grating orientations), participants’ responses on the current trial are subject to biases in an interceptive task (e.g., Kwon and Knill, 2013; de Lussanett et al. 2001). If such confoundings are not dealt with before looking at serial biases, these could lead to spurious serial effects (Pascucci et al. 2019). In the present work, we chose to use a multiple linear regression framework where interpretation of the previous trial speed slope relies on controlling for the effect of the current trial, and the results of this approach are similar to the process of residualization used elsewhere (Pascucci et al. 2019; Bliss et al. 2017).

For temporal errors, negative values for the previous trial speed regression slopes indicate an attractive serial bias toward previous trial information. In other words, the greater the speed of the previous trial, the more participants tend to anticipate their responses. It is important to note that, in the present experiment, we cannot know whether participants store information about previous trial speed, time-to-contact, or motor response. Despite not being able to dissociate these three features of the task in our study, all three are correlated and should not change our interpretation of the results (e.g. higher speeds lead to shorter time-to-contacts and faster responses on the current trial). To illustrate that a negative slope for the previous trial speed regressor represents an attractive bias toward previous trial speed (or time-to-contact or motor response), we will give an example assuming the bias is toward the previous trial speed. Take a current trial where the target travels at 24 dva/s and that was preceded by a trial with a target traveling at 28 dva/s. In this case, an attractive bias would be represented by the current speed being processed and perceived as being more similar to the previous trial speed than it actually is. Let us assume this biased representation of speed takes the value of 25 dva/s. If the current target is traveling at a speed of 24 dva/s and the brain represents it as a target speed of 25 dva/s and assuming the participant’s response is guided by such speed, the response to the current target will be a little bit faster than it should. For a traveling distance of 10 dva, the target will take 416 ms to arrive at the interception point if it is traveling at 24 dva/s, whereas for a target traveling at 25 dva/s it would take 400 ms. In that situation, the participant would anticipate their responses in 16 ms.

To evaluate the presence of serial dependence for each ITI, we performed one-tailed one-sample t-tests on regression slopes. To evaluate the serial dependence dynamics over ITIs, previous trial slopes for each ITI were entered in a one-way repeated-measures ANOVA. A Helmert contrast was used to follow up on significant ANOVA results (alpha = 0.05). The analysis was performed on Python and statistical analysis was performed on JASP (0.13.1.0).

To check whether the serial dependence effect was not due to spurious correlations present in the particular trial sequence presented to each participant, we regressed the current trial error with the next trial speed, expecting the absence of an effect (Bliss et al. 2017; Fischer and Whitney, 2014). If temporal errors on the current trial were found to be affected by the next trial target speed, this would suggest that spurious correlations are present in the particular trial sequence presented to each participant. Finding such association with the next trial would make one question the effects with previous trial information (Pascucci et al. 2019).

## Results

The goal of the present experiment was to verify whether increasing the time between trials would decrease the serial dependence effect in a coincident timing task. In such a task, after accounting for current trial biases, an attractive bias toward previous trial speed (or toward previous trial time-to-contact or motor responses, as these information are correlated in our task) would be evidenced by responses getting more anticipated as previous trial speeds increase.

Our results showed a statistically significant serial effect for all ITIs, as evidenced by a negative slope for the previous trial speed regressor (Table 1). As shown in Figure 2A, average temporal error decreases as previous trial speed (x axis) increases for each current trial speed (vertical axis), indicating an attractive bias toward the previous trial. To illustrate this bias in more detail, take for example when the current speed is 24 dva/s (middle row of Figure 1D, left panel). In this case, when the previous speed was 28 dva/s, the current speed (or time-to-contact or motor response, which are correlated in our current design) might be perceived as being faster than it actually is and this would lead to an anticipated response (as can be seen in the rows of Figure 1D (left panel) and Figure 2A). This bias occurs for all current trial speeds but at different ranges of temporal error (as faster current trial speeds induced later responses, which will be later discussed). This attractive bias is in line with previous studies in interceptive actions (Kwon and Knill, 2013; Makin et al. 2008) and is present in all ITI conditions (Figure 2B left).

**Figure 2.**
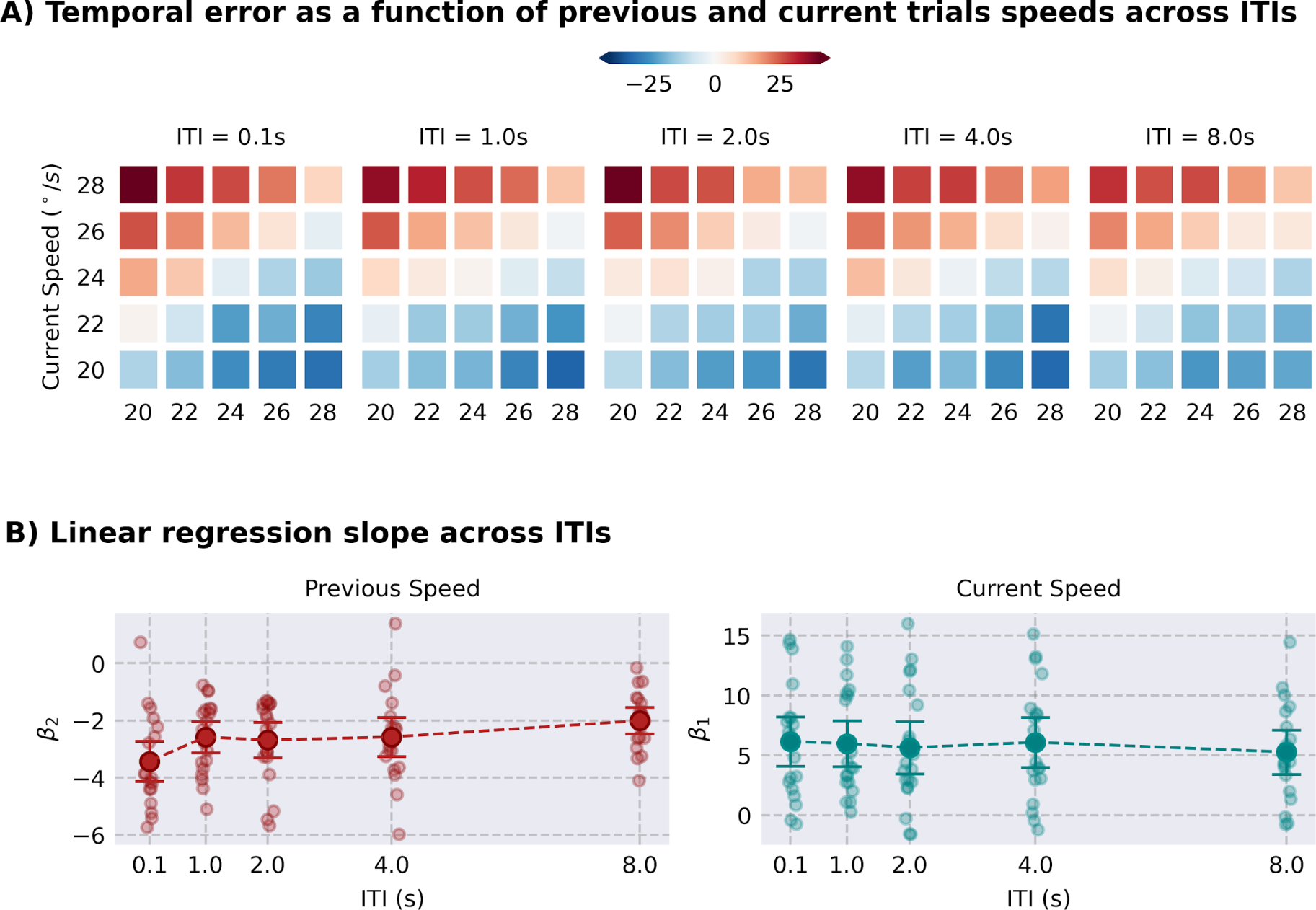
Serial dependence abruptly decreases after 0.1 s ITI but stabilizes afterward. (A) Attractive serial bias pattern of the temporal error as a function of current (vertical axis) and previous (horizontal axis) trial speed for all ITIs. The attractive serial bias is evidenced by temporal errors becoming more negative as previous trial speed increases. (B) Previous (left) and current (right) speed slopes of multiple linear regressions performed on individual participant data for all ITIs. The closer a value is to zero, the lower the regressor's contribution for explaining the temporal error. Large opaque dots represent average parameter estimates across participants, small and transparent dots represent individual data, and error bars represent 95% confidence intervals over parameter estimates.

**Table 1.** Statistical results of the one-sample one-tailed t-tests used to verify the presence of serial dependence for each ITI on temporal error.

| ITI | t(19) | p-value | Cohen's *d* |
| --- | --- | --- | --- |
| 0.1 s | −9.632 | < 0.001 | −2.154 |
| 1 s | −9.229 | < 0.001 | −2.064 |
| 2 s | −8.472 | < 0.001 | −1.894 |
| 4 s | −7.362 | < 0.001 | −1.646 |
| 8 s | −8.593 | < 0.001 | −1.971 |

When comparing serial dependence across ITIs, we found it decreased with increasing time between trials [F(4,72) = 3.963, p = 0.006, ω^2^ = 0.074], but that there was an abrupt decrease from 0.1 to 1 s [Helmert contrast between 0.1 s against all other ITIs: t(18) = –3.571, p <0.001], which then did not decrease between 1 s to 8 s ITIs [all p>0.05 for subsequent contrasts] (Figure 2B left).

There was also a statistically significant effect of current trial speed for all ITIs (Table 2), as evidenced by positive slopes of the current trial speed regressors. In addition, current trial speed regression slope was not modulated by ITIs [F(4,72) = 0.469, p = 0.758, ω^2^ = 0.000]. Note that the effect of the current trial speed is opposite to that of the previous trial speed on temporal error: the greater the speed of the target on the current trial, the more participants tend to present delayed responses with respect to the target (Figure 1C and Figure 2A and 2B right panel). This does not mean, however, that participants’ responses are repulsed away from the previous trial speed. As explained above, if participants’ responses are attracted toward previous trial speed (or time-to-contact or motor response), we should expect more anticipated responses as previous trial target speed gets faster, after controlling for the effects of the current trial. On the other hand, faster targets on the current trial elicit more delayed responses compared to slower targets because participants’ responses are not as fast as they should be, but still their absolute response time is faster than for the slower targets. In addition, the current trial speed bias might emerge due to a central tendency effect (e.g. Jazayeri and Shadlen, 2010). Central tendency alone, however, cannot explain the bias caused by the previous trial speed (Kwon and Knill et al. 2013; Motola et al. 2020).

**Table 2.** Statistical results of the one-sample one-tailed t-tests used to verify the presence of current trial biases for each ITI on temporal error.

| ITI | t(19) | p-value | Cohen's *d* |
| --- | --- | --- | --- |
| 0.1 s | 5.831 | < 0.001 | 1.304 |
| 1 s | 6.154 | < 0.001 | 1.376 |
| 2 s | 5.029 | < 0.001 | 1.125 |
| 4 s | 5.695 | < 0.001 | 1.273 |
| 8 s | 5.587 | < 0.001 | 1.282 |

Given that the serial dependence effect was pronounced even after 8 s, we next asked how stable the memory trace that leads to serial dependence might be. According to theoretical models of short-term memory, long delays lead to higher response variability in short-term memory tasks due to memory trace diffusion (e.g., Schneegans and Bays 2018; Wimmer et al. 2014). If this is the case, we should see an increase in error variability as a function of ITI, allowing for the memory trace to diffuse more. To characterize how and whether variability increased as a function of the ITI, in a first step, we averaged across current and previous speeds and compared error variability over ITIs. Although we find evidence for a change in error variability over ITIs [F(4, 72) = 5.902, p < 0.001, ω^2^ = 0.029], this was due to a decrease of error variability from 0.1 s compared to the rest of ITIs [Helmert contrast between 0.1 s against all other ITIs: t(18) = 4, p <0.001] and from 4 s compared to 8 s [Helmert contrast between 4 s and 8 s ITIs: t(18) = 2.727, p = 0.008] (Figure 1B [width of response distribution] and 3B).

To explore further how different ITIs modulate response variability, we performed a similar multiple regression on error variability as before, checking the dynamics of error variability across ITIs as a function of previous and current trial speed for all participants and ITIs. Our results show that error variability decreased as target speed increased in the current trial, as evidenced by a positive slope in the multiple regression for all ITIs (Table 3 and Figure 3B right), and that error variability increased as target speed increased on the previous trial for all ITIs, except 4 s (Table 4 and Figure 3B right). In addition, we found no evidence for a change for both current [F(4,72) = 0.885, p = 0.478] and previous [F(4,72) = 0.274, p=0.894] trial speed regression slopes across ITIs (Figures 3A and 3B).

**Figure 3.**
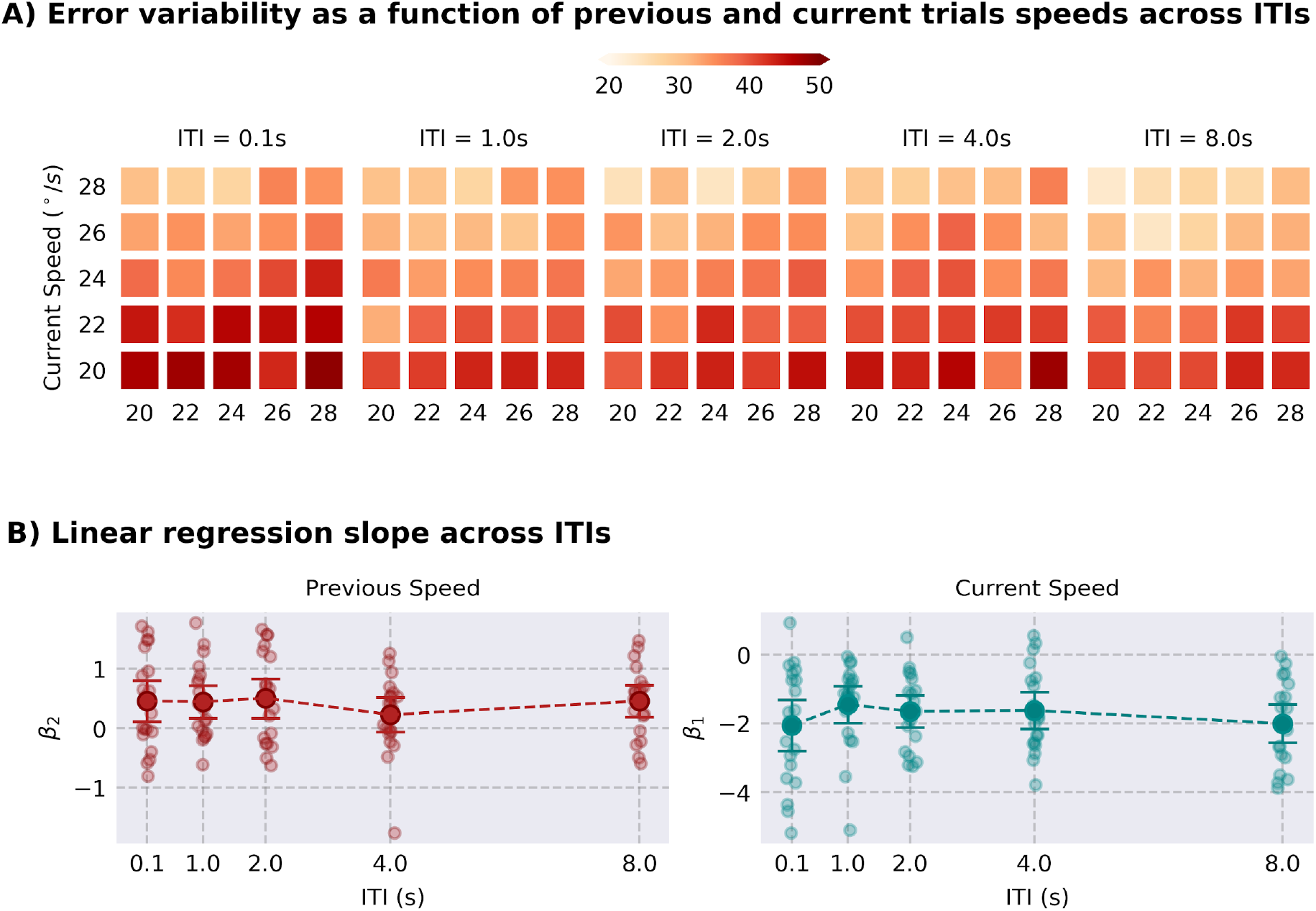
Error variability decreases with increased current target speed and is stable across ITIs. (A) Error variability as a function of current (vertical axis) and previous (horizontal axis) trial speed. There is a tendency for error variability to decrease as current trial speed increases (lighter colors from bottom to top) and increase as previous trial speed increases (darker colors from left to right). (B) Previous (left) and current (right) slopes of multiple linear regressions performed on individual participant error variability across ITIs. The closer a value is to zero, the lower the regressor’s contribution for explaining the error variability. Large opaque dots represent average parameter estimates across participants, small and transparent dots represent individual data, and error bars represent 95% confidence intervals over parameter estimates.

**Table 3.** Statistical results of the one-sample two-tailed t-tests used to verify the presence of current trial biases for each ITI on response variability.

| ITI | t(19) | p-value | Cohen's *d* |
| --- | --- | --- | --- |
| 0.1 s | −5.454 | < 0.001 | −1.220 |
| 1 s | −5.309 | < 0.001 | −1.187 |
| 2 s | −6.911 | < 0.001 | −1.545 |
| 4 s | −5.980 | < 0.001 | −1.337 |
| 8 s | −7.008 | < 0.001 | −1.608 |

**Table 4.** Statistical results of the one-sample two-tailed t-tests used to verify the presence of previous trial biases for each ITI on response variability.

| ITI | t(19) | p-value | Cohen's *d* |
| --- | --- | --- | --- |
| 0.1 s | 2.557 | 0.019 | 0.572 |
| 1 s | 3.188 | 0.005 | 0.713 |
| 2 s | 2.997 | 0.007 | 0.670 |
| 4 s | 1.524 | 0.144 | 0.341 |
| 8 s | 3.328 | 0.004 | 0.763 |

We also performed a control analysis to check whether participants were performing the task correctly and whether serial dependence results were not due to spurious patterns on the data. Merely reactive responses would be indicated by temporal errors on the order of 200 ms. Participants’ temporal errors were lower than 200 ms (mean temporal error (lower CI, upper CI) = –72.13 (−96.72, –47.54); t(19) = −21.689, p<0.001), indicating that they were indeed using the time to contact to perform the task rather than waiting for the target to hit the vertical bar (Figure 1B). As another control analysis, we should not expect that temporal errors on the current trial be affected by the next trial target speed as trial-to-trial target speeds are independent. Performing a regression analysis of current trial temporal errors by the next trial speed showed absence of evidence for this effect over all ITIs (ITI = 0.1s: t(19) = 0.357, p = 0.712; ITI = 1 s: t(19) = 1.569, p = 0.133; ITI = 2 s: t(19) = 1.345, p = 0.195; ITI = 4 s: t(19) = 1.435, p = 0.168; ITI = 8 s: t(19) = 0.527, p = 0.605), indicating that the effect caused by the previous trial speed is not due to spurious correlations in the data.

## Discussion

How long does the serial dependence effect persist between trials? Previous studies have shown that the impact of a trial on future behavior decreases with time (Fischer and Whitney, 2014; Kalm and Norris 2018; Fritsche et al. 2020). However, this decrease in serial dependence might be explained by an overwriting of the representation of previously experienced stimuli or responses (Matthey, Bays and Dayan 2015) or by decreasing their representation precision due to interference (Kalm and Norris 2018), not temporal decay alone. To understand the sole effect of the passage of time into the memory trace that underlies serial dependence, we had participants perform a coincident timing task at different inter-trial intervals in separate blocks. Our results indicate that the serial dependence effect fades abruptly from 0.1 s to 1 s, but from there on remains pronounced for up to 8 s. In addition, participants’ response variability seemed to be stable even after long intervals between trials, as evidenced by decreased response variability for longer ITIs. These results indicate that the memory trace that causes serial dependence is stable over at least 8 s.

Previous studies have also investigated the temporal properties of serial dependence, although with substantial differences in results. In Bliss et al. (2017), serial dependence was tested in a delayed reproduction task. The authors found that serial dependence sharply dropped from 3 to 6 s, and for intervals of 10 s, it turned into a repulsive bias. Conversely, and consistent with the results of the present work, Papadimitrou et al. (2015) showed that for 2 out of 3 monkeys, serial dependence was still high after 6 s of inter-trial interval, and that, similar to Bliss et al. (2017), one of the monkeys had serial dependence drop to zero after 6 s. A possible explanation for these contradictory results could be found in task differences. For instance, Bliss et al. (2017) used a delayed visual reproduction task, which might rely more heavily on visual memory than our coincident timing task and Papadimitrou et al. (2015) delayed saccade task. Another critical difference is that, in our task, participants were not required to keep target information actively in memory. In contrast, in Bliss et al. (2017) and Papadimitrou et al. (2015), keeping current trial information in working memory was required to perform the task.

It might be the case that implicitly and explicitly encoded memory traces have different temporal dynamics. There is evidence that working memory representation loses precision over longer delay intervals (Rademaker et al. 2018; Shin et al. 2017). These results have been interpreted as an increase in the uncertainty of memory representation over time (c.f. Pouget et al. 2013). Assuming memory information is combined with current sensory information weighted by their uncertainty to produce serial dependence (van Bergen and Jehee, 2019; Fritsche et al. 2020), a steady decrease of the memory trace precision should have led to a steady reduction of serial dependence in our experiment. However, we observed an abrupt decrease in response bias for short ITIs (1 s), from where it is still present for up to 8 s. In contrast to previous working memory studies (Rademaker et al. 2018; Shin et al. 2017), these results suggest that the memory trace’s precision implicitly acquired in a coincident timing task does not vanish quickly in the absence of interference.

Our results show an abrupt change in serial dependence from 0.1 s to 1 s of waiting time between trials. One possibility to explain such an abrupt change and later stabilization is that the information from previous trial changes in its storage mode, modulating how uncertainty is represented. It could be that, at 0.1 s, there is still activity associated with the ongoing processing of the previous trial, whereas at 1 s the information about the last trial is already stored in short-term memory. In this case, when changing from a perceptual/motor activity into a short-term memory representation, uncertainty might jump to a higher level and stay stable over time. An alternative (and somewhat simpler) explanation to such a result is that the abrupt decrease in serial dependence is associated with the time constant of the decay of uncertainty information in short-term memory. Although there is evidence showing that stimulus uncertainty is kept in working memory (Honig et al. 2021) and that perceived variance on the current trial is affected by the perceived variance on the previous trial (Suárez-Pinilla et al. 2018), it is also possible that the uncertainty associated with the information that leads to serial dependence is related to the previous trial decision or response. In support of this view, it has been shown that higher confidence levels on the previous trial response increase the serial dependence effect even when the previous trial stimulus uncertainty is the same (Samaha et al. 2019). In addition, manipulating the uncertainty of the stimulus in the previous trial does not affect serial dependence (Ceylan et al. 2021). This is an interesting topic and it remains an open question how much the uncertainty stored in implicit memory relies on the uncertainty of the previous trial sensory information.

It has been suggested that serial dependence could result from a reminiscent and persistent activity encoding the previous target until the subsequent trial (Papadimitrou et al. 2015). Assuming that such a proposal is true, current attractor network models used to explain short-term memory storage would predict that the longer the delay, the greater the variability in participants’ responses (e.g., Wimmer et al. 2014). Given that we observed a decrease in response variability for longer intervals between trials in our data, we speculate that this is due to a different information storage mode. Our findings are more consistent with recent models that have proposed that information could be stored in short-term memory through changes in synaptic weight (Mongilo et al. 2008). This mechanism would lead to a reduced diffusion of the memory content stored in the network (Itskov et al. 2011; Seeholzer et al. 2019), resulting in a temporally stable representation of memory contents. Our results also suggest that such an “activity-silent” mechanism should decrease the rate of decay of the memory trace. Although the idea of memory being stored in silent, hidden states has received electrophysiological evidence (Barbosa et al. 2020; Stokes 2015; Wolff et al. 2017; Rose et al. 2016), these results are far from unanimous (Papadimitrou et al. 2017), and it might be the case that the exact mechanism is task-dependent. However, it is essential to note that such models were exclusively created based on prefrontal neurons’ activity, and the memory trace that leads to serial dependence might be stored elsewhere (c.f. Christophel et al. 2017).

An alternative explanation of why we do not find evidence for an increase of error variability over time is a possible limitation of our task when compared to purely perceptual tasks: the motor component of the coincident timing task might overshadow the noise due to diffusion. It could be argued that noise present in generating the motor command and processing sensory information would be unsusceptible to the effect of time. However, if these two sources of noise exist and are larger than the impact of noise due to diffusion of the memory trace, then the effects of diffusion would be difficult to show in our data. Another possible explanation for a slight decrease in response variability as participants wait more time is that they get more attentive as they get more rested (although we would expect that waiting more time between trials might have made participants more distracted, which would have increased their response variability). Future studies using tasks that control other sources of noise might help to resolve this issue.

There is a growing interest in the serial dependence literature to identify the source of serial dependence: whether it comes from sensory signals (Fischer and Whitney 2014; Cicchini et al. 2017), biased percepts (Cicchini et al. 2021; Collins 2020; Fornaciai and Park 2020), decision processes (Ceylan et al. 2021; Fischer et al. 2020; Fritsche et al. 2017; Pascucci et al. 2019; Roseboom 2019) or motor responses (St. John Saaltink et al. 2016). A limitation of the present work is that speed, time-to-contact and motor responses are correlated, preventing us from pinpointing the source of serial dependence in the coincident timing task. Cicchini et al. (2017) have shown that response errors were not correlated with previous trial responses when these were orthogonal to previous stimulus orientation in a delayed reproduction task. However, since the coincident timing task has a strong reliance on motor action, this could have a greater influence on current trial responses and this should be an interesting topic to be addressed in future studies. Although we use speed as the independent variable in the multiple linear regression, the fact that the task attributes are correlated (high speeds lead to shorter time-to-contact and faster motor responses) does not change the interpretation of the current results that the bias from the previous trial changes as a function of ITI.

Our results provide empirical evidence that serial dependence is sustained over time and the memory trace underlying it is stable, suggesting that future studies and models should look for explanations that accommodate such slow decay and stability. We speculate that the memory trace that leads to serial dependence is different from the one proposed to keep information through a persistent activity during the delay period in working memory tasks and favor the hypothesis that implicit short-term memory is stored through short-term synaptic plasticity mechanisms.

### Open practices statement

Task and analysis code and raw data will be made openly available, following acceptance of the manuscript at Open Science Framework (https://osf.io/8qdnw/?view_only=51e2287fa5084a29b48e8f15bcd149ef). The procedures and analysis were not pre-registered prior to the research being conducted.

